# Cocaine-Induced Impulsivity is Differentially Expressed in Male and Female Mice Exposed to Maternal Separation and is Associated with Alterations in AMPA Receptors Subunits

**DOI:** 10.1101/2020.06.05.136812

**Authors:** Adriana Castro-Zavala, Ana Martin-Sanchez, Larisa Montalvo-Martínez, Alberto Camacho-Morales, Olga Valverde

**Affiliations:** Neurobiology of Behaviour Research Group (GReNeC-NeuroBio), Department of Experimental and Health Sciences, Universitat Pompeu Fabra, Barcelona, Spain; Department of Biochemistry, College of Medicine, Universidad Autónoma de Nuevo León, Monterrey, C.P. 64460, México; Neurometabolism Unit. Center for Research and Development in Health Sciences, Universidad Autónoma de Nuevo León, Monterrey, C.P. 64460, México; Neuroscience Research Program, IMIM-Hospital del Mar Research Institute, Barcelona, Spain

**Author notes:** Corresponding author: MD. PhD. Olga Valverde, Neurobiology of Behaviour Research Group (GReNeC - NeuroBio), Department of Experimental and Health Sciences, Universitat Pompeu Fabra, Dr. Aiguader 88; Barcelona 08003.

**Keywords:** cocaine self-administration, depression-like behaviour, Gria1, Gria2, GluA1, GluA2

## Abstract

Impulsivity is a key trait in the diagnosis of major depressive disorder (MDD) and substance use disorder (SUD). MDD is a chronic illness characterized by sadness, insomnia, and loss of interest. SUD is a chronic and relapsing disorder characterized by the consumption of drugs despite their negative consequences. Among drugs of abuse, cocaine is the most consumed psychostimulant. Animal studies demonstrated that increased impulsivity predicts predisposition to acquire cocaine self-administration (SA) behaviour with an increased cocaine-intake. Moreover, early-life stress represents a vulnerability factor to develop depressive disorders and drug addiction. Maternal separation with early weaning (MSEW) is an animal model that allows examining the impact of early-life stress on cocaine abuse. In this study, we aimed to explore changes in MSEW-induced impulsivity to determine potential associations between depression-like and cocaine-seeking behaviours in male and female mice. We also evaluated possible alterations in the AMPA receptors (AMPArs) composition and glutamatergic neurotransmission. We exposed mice to MSEW and the behavioural tests were performed during adulthood. Moreover, GluA1, GluA2 mRNA and protein expression were evaluated in the medial Prefrontal Cortex (mPFC). Results showed higher impulsive cocaine-seeking in females, independently the MSEW, as well as an increase in GluA1 and GluA2 protein expression. Moreover, MSEW induced downregulation of Gria2 and increased the GluA1/GluA2 ratio, only in male mice. In conclusion, female mice expressed higher mPFC glutamatergic function, which potentiated their impulsivity during cocaine SA. Also, data indicated that MSEW alters glutamatergic function in mPFC of male mice, increasing the glutamatergic excitability.

## INTRODUCTION

Impulsivity is a major personality and temperament dimension consisting of maladaptive behaviour and characterized by poorly conceived, prematurely expressed, unduly risky, or inappropriate actions often resulting in undesirable consequences (Granö *et al.* 2007; Dent and Isles 2014; Dalley and Ersche 2019). Therefore, this personality trait can increase predisposition to suffer drug addiction (Nicholls *et al.* 2014; Rømer Thomsen *et al.* 2018; Adams *et al.* 2019; Butelman *et al.* 2019; Jupp *et al.* 2020) and other psychiatric disorders like major depressive disorder (MDD) (Dent and Isles 2014; Dalley and Ersche 2019). Depression is the most common psychological disorder affecting more than 264 million people worldwide (James *et al.* 2018). This disorder is characterized by sadness, loss of interest or pleasure, feelings of guilt, low self-worth, disturbed sleep or appetite, tiredness, and poor concentration (WHO 2017),

Impulsivity has been also proposed as an endophenotype to investigate the underlying neurobiological mechanisms for many impulse control disorders including substance use disorder (SUD) (Dalley and Ersche 2019). Among illicit drugs, cocaine is one of the most consumed psychostimulants with more than 18 million users (UNODC 2019). Clinical studies show that cocaine addiction includes poor inhibitory control for goal-directed behaviour in the frontal cortical regions, evidencing an impulse control disorder which induces drug craving (Barrós-Loscertales *et al.* 2019). In line with this, preclinical studies demonstrated that increased impulsivity predicts predisposition for drug use to evolve towards drug abuse (Jupp *et al.* 2020).

Animal studies show that rats with high impulsivity met the acquisition criterion faster, in greater percentage and also consumed more cocaine than rats with low impulsivity (Perry *et al.* 2005). In females rats, it was observed that impulsivity could predict the levels of acquisition of cocaine self-administration (SA) (Perry *et al.* 2005). Moreover, in male rats, high impulsivity predicts the development of compulsive cocaine use (Belin *et al.* 2008). Additionally, in a genetic animal model of impulsivity, Roman high-avoidance rats showed increased cocaine sensitization (Giorgi *et al.* 2005), higher number of responses in the SA paradigm (Fattore *et al.* 2009) and higher vulnerability to self-administer cocaine (Fattore *et al.* 2009). All of these behavioural alterations were associated with decreased volume and function of some mesocorticolimbic areas related with the development and persistence of cocaine addiction (Fattore *et al.* 2009; Giorgi *et al.* 2019; Río-Álamos *et al.* 2019).

Several studies have also demonstrated that cocaine exposure induces changes in the glutamatergic system, including the AMPA receptors (AMPArs) subunit composition (Bowers *et al.* 2010; Castro-Zavala *et al.* 2020a; Castro-Zavala *et al.* 2020b). AMPArs are made up of four subunit proteins (GluA1-A4) and generally composed of GluA2 in complex with GluA1 or GluA3 (Bowers *et al.* 2010). Therefore, previous works showed increased AMPArs function after cocaine exposure, because of the induction of long-term potentiation (Kauer and Malenka 2007). This high activity of AMPArs could be explained by the insertion of GluA2-lacking AMPArs, because this kind of receptors are calcium-permeable, have greater channel conductance and trigger calcium-dependent signalling cascades (Bowers *et al.* 2010). Regarding impulsivity and the vulnerability to cocaine abuse, Nakamura *et al* (2000) showed that the administration of NBQX, an AMPArs antagonist, decreased impulsivity in a dose-dependently way in rats. Additionally, Barkus *et al.* (2012) making use of a GluA1 knock-out mice model with an impulsive phenotype, reported a faster acquisition in the food SA paradigm and a decreased capacity to extinguish the SA behaviour. These studies evidenced the key role played by the AMPArs in the modulation of impulsivity.

As previously mentioned, there is also a close relationship between impulsivity and MDD. In fact, GluA2-lacking AMPArs were suggested to be a common link between depression and SUD (Goffer *et al.* 2013; Martínez-Rivera *et al.* 2017; Castro-Zavala *et al.* 2020b). Clinical studies reported increased impulsivity in adults with a later MDD diagnosis, suggesting impulsivity as a predictor of the development of major depression (Granö *et al.* 2007). Also, Corruble *et al.* (2003) described three characteristics of impulsivity in adults with severe depression: behavioural loss of control, non-planned activities and cognitive impulsivity. In line with these studies, some authors have evaluated the association between impulsivity and childhood adversity in depressed adults (Brodsky *et al.* 2001). These studies reported that those patients with history of childhood trauma showed higher impulsivity and higher suicidal behaviour than the ones without childhood adversity (Brodsky *et al.* 2001).

Maternal separation with early weaning (MSEW) is an animal model that allows researchers to reproduce the effects of childhood adversity(George *et al.* 2010; Vetulani 2013; Bian *et al.* 2015). Moreover, MSEW induces a depression phenotype that permits the examination of the impact of early-life stress on cocaine use or abuse (Liu *et al.* 2018; Vannan *et al.* 2018; Castro-Zavala et al., 2020b). We have previously reported, using a model of MSEW, an increased depression- and anxiety-like behaviour(Gracia-Rubio *et al.* 2016b; Portero-Tresserra *et al.* 2018), higher acquisition in the cocaine SA paradigm in males but not in females (Castro-Zavala *et al.* 2020b) and increased GluA1/GluA2 in the NAc and VTA of males exposed to MSEW and cocaine SA (Castro-Zavala *et al.* 2020b). Moreover, we also observed an increased acquisition percentage in females, which was independent of the early-life stress (Castro-Zavala *et al.* 2020b), being in accordance with the telescoping effect observed in women regarding drug use disorders (Haas and Peters 2000).

In this context, this work aimed to explore the changes in impulsivity induced by the MSEW model in males and females CD1 mice. Moreover, we determined a possible association between depression-like behaviour, impulsive cocaine-seeking, as well as alterations of AMPArs subunit (mRNA and protein levels). To achieve these goals, we evaluated impulsivity for cocaine intake (as a percentage of response efficiency) in the SA paradigm. We also determined GluA1 and GluA2 protein expression and evaluated Gria1 and Gria2 mRNA expression in the medial prefrontal cortex (mPFC), a brain area involved in inhibitory control of behaviours.

## MATERIALS AND METHODS

### Animals

Sixteen male and sixteen female CD1 adult mice of 10 weeks of age were used as breeders (Charles River, Barcelona, Spain). Animals were received at our animal facility, UBIOMEX, PRBB. The animals were placed in pairs in standard cages at a temperature- (21 ± 1ºC) and humidity- (55% ± 10%) controlled room and subjected to a 12 h light/dark cycle; with the lights on from 8:00 to 20:00 h and *ad libitum* access to food and water. Ten days later, the males were removed from the cages. Once offspring had been weaned, mice were assigned randomly to the cocaine SA, food SA or to naïve condition. A different group of mice was used to perform the tail suspension test (TST). The total number of animals used for this work was 189. The experiments were carried out in accordance with the guidelines of the European Communities Directive 88/609/EEC regulating animal research. The local ethical committee (CEEA-PRBB) approved all procedures, and every effort was made to minimize animal suffering and discomfort as well as the number of animals used.

### Rearing conditions

The rearing conditions were as previously described (Gracia-Rubio *et al.* 2016b; Portero-Tresserra *et al.* 2018; Castro-Zavala *et al.* 2020a; Castro-Zavala *et al.* 2020b). New-born mice were randomly assigned to the experimental groups: standard nest (SN) and MSEW (Figure 1A). The day of birth was considered the postnatal day (PD) 0. Animals in the MSEW were separated from their mothers for 4 h per day (9:00 to 13:00 h) from PD2 to PD5, and 8 h per day (9:00 to 17:00 h) from PD6 to PD16. As for the separation, the mother was removed and placed in another cage and room, leaving the pups in their home box. To maintain the body temperature of the pups, home boxes were placed upon electric blankets until the mother was duly returned. Animals in the SN remained with their mother until weaning (PD21), whilst animals in the MSEW were weaned at PD17. In both cases (SN and MSEW), cages were cleaned on PD10. We distributed the pups of each litter between the different experimental groups in order to avoid a litter effect. MSEW procedure does not affect body weight (Gracia-Rubio *et al.* 2016a; Portero-Tresserra *et al.* 2018), mortality (George *et al.* 2010), morbidity (George *et al.* 2010) or the male/female ratio (Koob and Zorrilla 2010).

**Figure 1.**
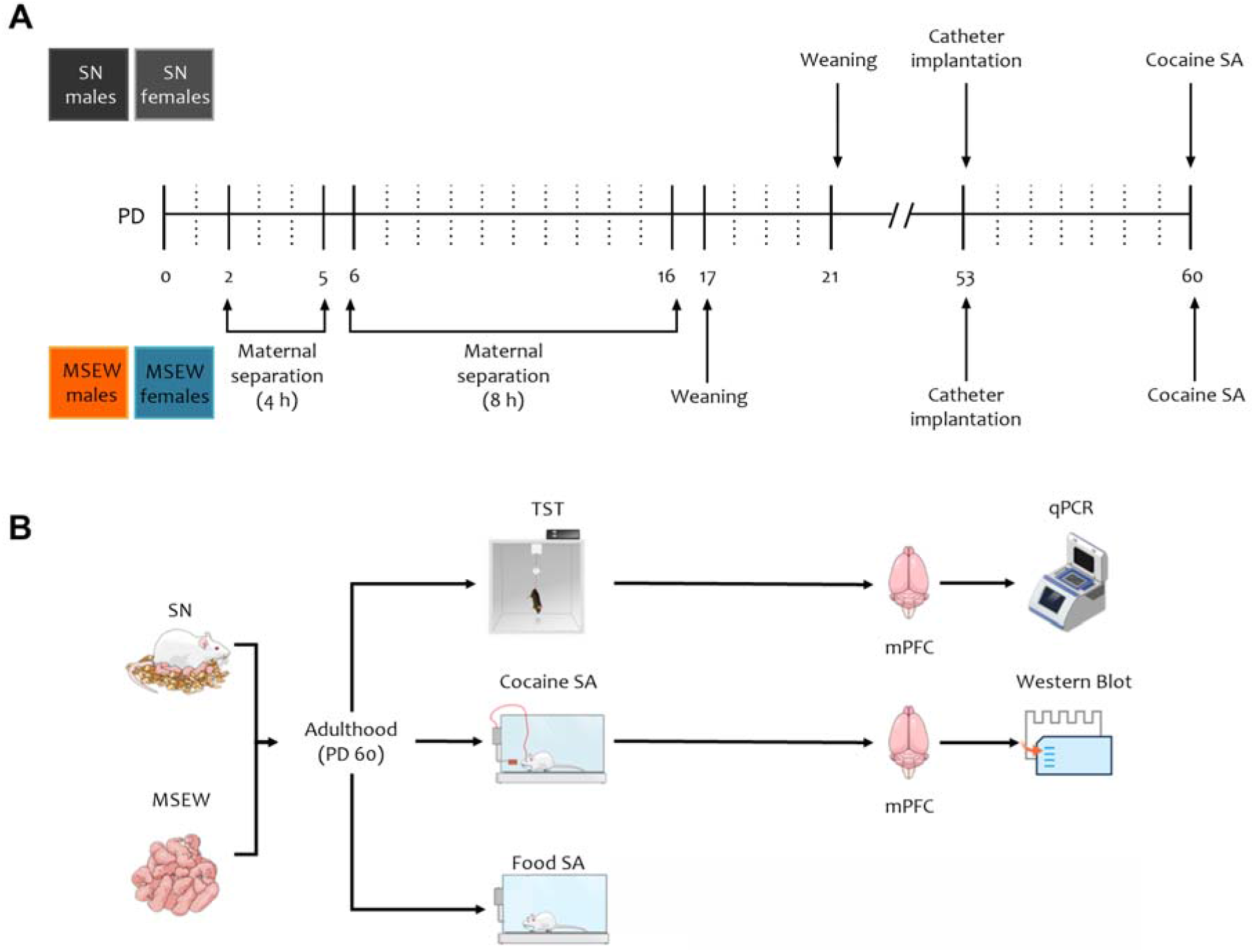
Schematic representation of the experimental schedule. (A) Schematic representation of the MSEW model and the (B) timeline in which the experiments were performed.

### Tail suspension test

Mice underwent the TST on PD60 as previously described (Gracia-Rubio *et al.* 2016b). Briefly, each mouse was suspended 50 cm above a benchtop for 6 minutes (using adhesive tape attached 1cm from the tip of the tail). The time (s) that the animal was immobile during this interval was recorded.

### Drugs

Cocaine hydrochloride was purchased from Alcatel (Ministry of Health, Madrid, Spain) and was dissolved in sterile physiological saline (0.9%, NaCl solution). A dose of 1 mg/kg/infusion was used for the acquisition phase of the SA procedure.

### Apparatus for self-administration experiments

The SA experiments were carried out in mouse operant chambers (Model ENV-307A-CT, Medical Associates, Cibertec S.A., Madrid, Spain) containing two holes; one was defined as active and the other as inactive. Nose-poking into the active hole produced a reinforcement (cocaine infusion or a food pellet) that was paired with two stimulus lights, one of which was placed inside the nose-poke and the other above the active hole. Mice received a maximum of 150 reinforcements, and each reinforcement was followed by a 15 s time-out period, in which no cocaine infusions were delivered. Nose-poking into the inactive hole had no consequences. The side on which the active/inactive hole was placed was counterbalanced.

At the beginning of each session, the house light was ON for 3 s and OFF for the rest of the experiment. The session started with a food pellet release or a cocaine priming injection and 4 s presentation of the light cue, situated above the active hole.

### Cocaine self-administration

The SA experiments were conducted as described (Ferrer-Pérez *et al.* 2019; Castro-Zavala *et al.* 2020a; Castro-Zavala et al., 2020b). Briefly, when the SN (males n=23, females n=23) and MSEW (males n=30, males n=20) animals reached PD53, a jugular-vein catheter implantation was performed. The surgery was done following anaesthetization with a mixture of ketamine/xylazine (50☐mg/mL, 10 mg/mL, administrated in a volume of 0.15 mL/10g). Animals were treated with analgesic (Meloxicam 0.5 mg/kg; i.p, administrated in a volume of 0.10 mL/10g) and antibiotic solution (Enrofloxacin 7.5 mg/kg, i.p., administrated in a volume of 0.03 mL/10 g). After surgery, animals were housed individually, placed over electric blankets, and allowed to recover. At least 3 days after surgery, animals were trained, on a fixed ratio 1, to self-administer cocaine (1.0 mg/kg per infusion). During 10-day sessions (2 h each), the amount of nosepokes in the active- and the inactive hole (responses during time in and time out) and the infusion number (responses during time in) were counted. Mice were considered to have acquired a stable SA behaviour when the following criteria were met on 2 consecutive days: ≥5 responses in the active hole and ≥ 65% of responses in the active hole. All animals accomplished the 10 sessions independently of the day of acquisition.

### Food self-administration

Four days before testing commenced, mice were food-restricted and for that, mice were fed accordingly to the 95% of their body mass daily. Food restriction lasted the duration of food-maintained operant behaviour. Water was available *ad libitum* during the experimental phase. The animals were trained, on a fixed ratio 1, to nosepoke for food pellets (Grain-Based Rodent #5001, Test Diet, Sawbridgeworth, UK) for 10-day sessions (2 h each). The nosepoke number in the active- and the inactive hole (responses during time in and time out) and the pellet number (responses during time in) were counted.

### Percentage of response efficiency and motor impulsivity

The evaluation of motor impulsivity could be divided into two processes: impulsive action and impulsive choice (Dalley *et al.* 2011; Dalley and Ersche 2019). Impulsive action is measured by the incapacity to self-restraint and perform anticipatory actions, known as a failure of motor inhibition (Dalley *et al.* 2011; Dalley and Ersche 2019).

Adapting the formula employed by Hynes *et al.* (2018) to our SA paradigm, we calculated the percentage of response efficiency as an indirect measure of impulsivity. A 100% response efficiency represents enough responses to obtain the reinforcement (one food pellet/one nosepoke or one cocaine infusion/one nosepoke). Therefore, decreased response efficiency means an increased impulsive response.

For the cocaine SA, we used the following formula:

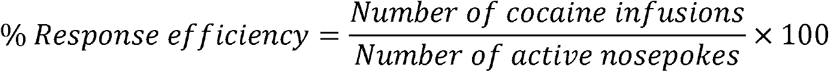

In the case of food SA, we calculated the percentage of response efficiency as follows:

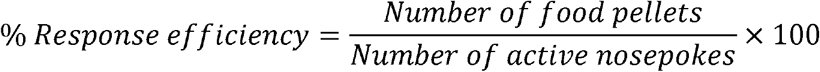

As a result, mice with high percentage response efficiency scores were considered less impulsive, while mice with low percentage response efficiency were considered more impulsive (Hynes *et al.* 2018).

### Animal Sacrifice and Sample Collection

Animals were sacrificed by cervical dislocation. Brains were immediately removed from the skull and placed in a cold plaque. Samples were dissected at different phases: without any behavioural test (drug-naïve), after cocaine SA (drug-experienced) and after TST (PD60). The drug-experienced animals are exclusively the mice that acquired the cocaine SA behaviour. mPFC was dissected and immediately stored at −80ºC until the biochemical analysis was performed. Samples from naïve- and drug-experienced mice were used for the western blot. For the qPCR, we utilised samples from mice that completed the TST.

### Western Blot for GluA1 and GluA2

To evaluate the expression of GluA1 and GluA2, samples were homogenized in cold lysis buffer (NaCl 0.15 M, EDTA 0.001 M, Tris pH 7.4 0.05 M, TX-100 1%, Glycerol 10%), supplemented with a protease inhibitor (Complete ULTRA Tablets Mini EASYpack, Roche, Mannheim, Germany) and a phosphatase inhibitor (PhosSTOP EASYpack, Roche, Mannheim, Germany). Protein samples (20 μg) were mixed with 5X loading buffer (TRIS pH 6.8 0.153 M, SDS 7.5%, Glycerol 40%, EDTA 5 mM, 2-β-mercaptoethanol 0.025%, bromophenol blue 0.025%), loaded and run on SDS-PAGE 10% and transferred to PVDF membranes (Millipore, Bedford, MA, USA). Membranes were blocked with BSA 5% for 1 h at room temperature and incubated overnight at 4ºC with primary antibodies (Table 1). Primary antibodies were detected with fluorescent secondary antibodies (Table 1), incubated for 1 h at room temperature. Images were acquired on a Licor Odyssey Scanner and quantified using Image Studio Lite software v5.2 (LICOR, USA). The expression of GluA1, GluA2 and β-tubulin were evaluated in the mPFC of the different groups: SN drug-naïve males, SN drug-experienced males, MSEW drug-naïve males, MSEW drug-experienced males, SN drug-naïve females, SN drug-experienced females, MSEW drug-naïve females and MSEW drug-experienced females (n=4-5 per group, run in triplicate). Data were normalized to the SN naïve males in order to determine the fold change due to sex, MSEW, cocaine exposure or the interaction between variables.

**Table 1.** Antibodies

| Antibody | # Catalogue | RRIDs | Dilution | Company |
| --- | --- | --- | --- | --- |
| GluA2 | AB1768 | AB_2313802 | 1:1000 | Milipore |
| GluA1 | ABN241 | AB_2721164 | 1:1000 | Milipore |
| β-tubulin | 556321 | AB_396360 | 1:5000 | BD Biosciences |
| goat anti-mouse IgG<br>H&L (IRDye 800CW) | ab216772 | AB_2857338 | 1:2500 | Abcam |
| goat anti-rabbit IgG<br>H&L (DyLight 680) | 611-144-002 | AB_1660962 | 1:2500 | Rockland |

### RNA isolation and real-time (RT)-PCR

RNA extraction from mPFC samples was performed using trizol as previously described (Cardenas-Perez *et al.* 2018). RT-PCR was performed by High-Capacity cDNA Reverse Transcription Kit (Applied Biosystems, Foster City, CA) using random primers and following standardized protocols.

### Quantitative PCR for Gria1 and Gria2

For the qPCR, we use cDNA (20 ng), Light Cycler SBYR green 480 Master Mix (Roche LifeScience, Product No. 04707516001) and the specific primers (Table 2) for Gria1, Gria2 and 36B4 as housekeeping gene (Integrated DNA Techologies, Inc.). The qPCR was performed in LightCycler ® 480 Instrument II (Roche LifeScience) using next program: 95°C-10s, 60°C-20s, 72°C-10s for 45 cycles.

**Table 2.**
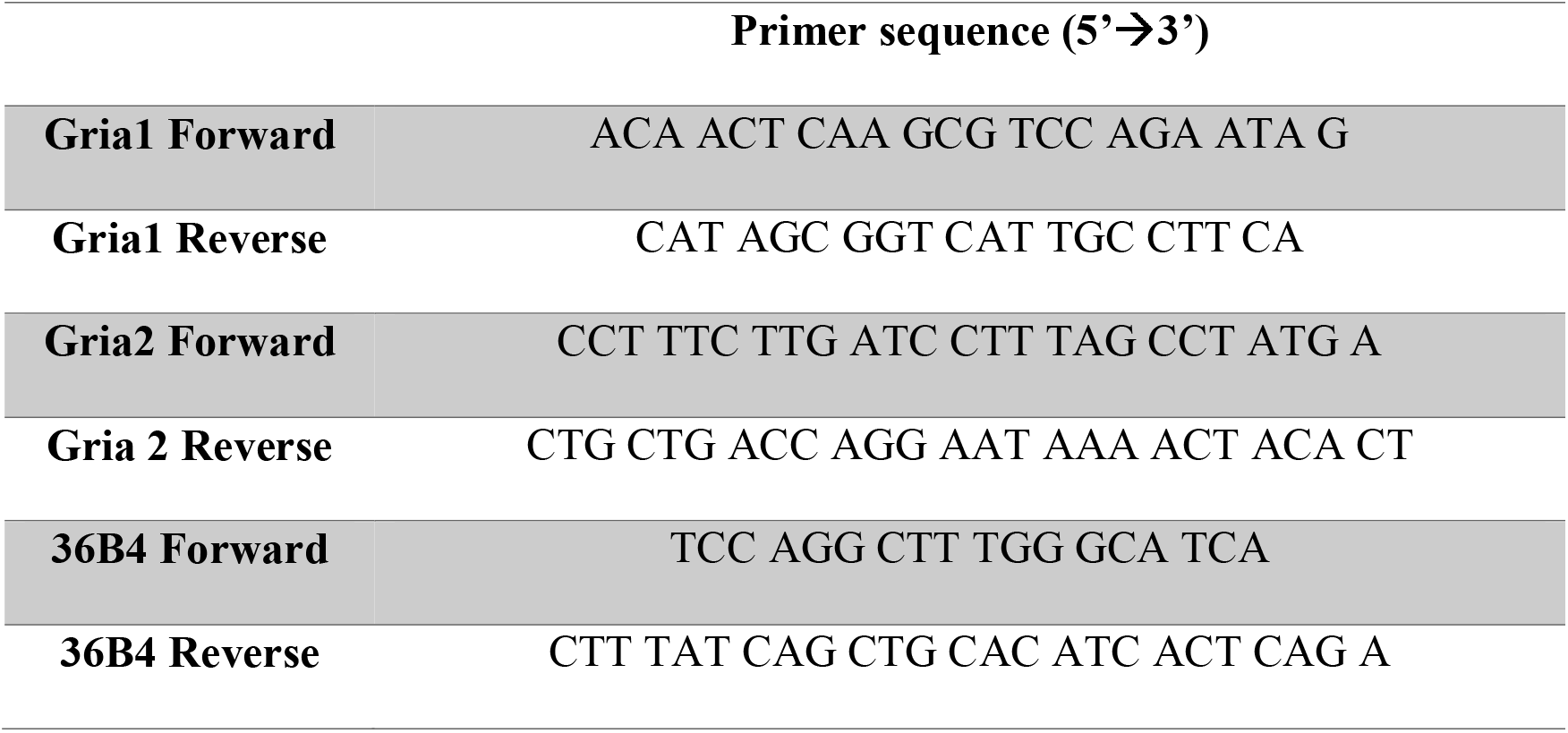
Primers for qPCR

### Statistical Analysis

Data were analysed for conditions of normality (Kolmogorov-Smirnov`s test), sphericity (Mauchly’s test) and homoscedasticity (Levene’s test). Data from the TST, qPCR, average of response efficiency and western blot results of drug-naïve mice were analysed using a two-way ANOVA with *rearing* and *sex* as independent factors. Western blot results of drug-naïve and drug-experienced animals were analysed using a three-way ANOVA with *rearing*, *sex* and *treatment* as factors. Data of percentage of response efficiency were analysed using a three-way ANOVA repeated measures with *rearing*, *sex* and *days* as factors of variation. When F achieved p<0.05, the ANOVA was followed by the Bonferroni post-hoc test if a main effect and/or interaction was observed. All possible post-hoc comparisons were evaluated. Statistical analyses were performed using SPSS Statistics v23. Data are expressed as mean ± SEM and a value of p<0.05 was considered significant.

## RESULTS

### Maternal separation increases depression-like behaviour only in male mice

The effects observed in the TST were evaluated in female and male mice during adulthood (PD60) in MSEW and control mice (Figure 2). A two-way ANOVA of immobility time showed a significant effect of *sex* (F_1,39_=12.324, p<0.01), *rearing* (F_1,39_=10.291, p<0.01) and the interaction between these factors (F_1,39_=4.908, p<0.05). The *sex* effect evidenced higher immobile time in females than males (p<0.01) while the *rearing* effect showed that MSEW mice remained immobile for a longer time (p<0.01). Bonferroni post-hoc test for the interaction showed that SN females spent more time immobile than SN males (p<0.001) and also that MSEW males showed a higher immobile time than SN males (p<0.01).

**Figure 2.**
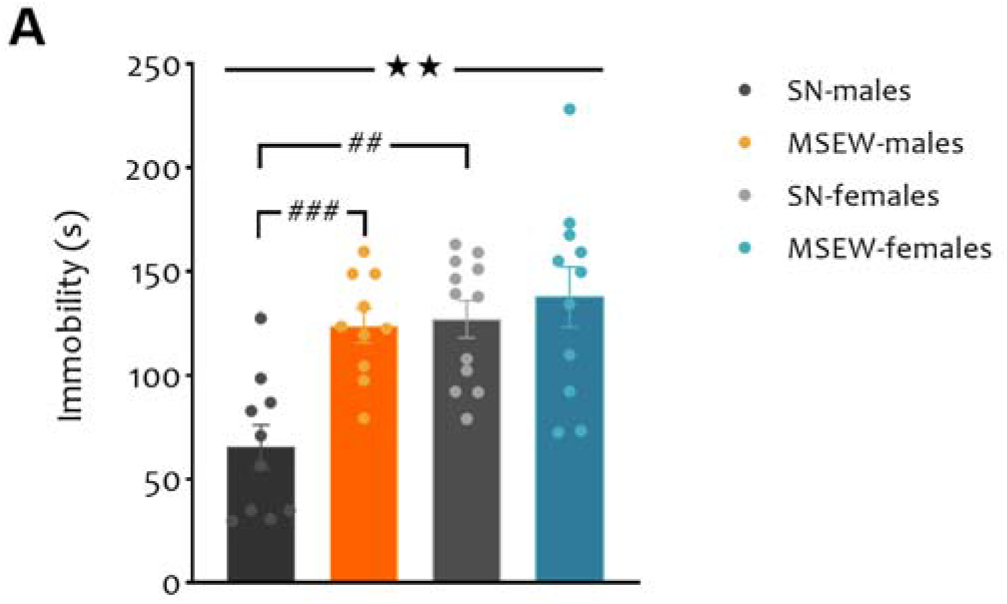
Effects of MSEW on the TST. Total immobility time in SN and MSEW adult male and female mice. *Sex* main effect of the ANOVA (⍰⍰p<0.01). Bonferroni post-hoc comparison for the interaction *sex* × *rearing* is indicated with the lines (##p<0.01, ###p<0.001). Data are expressed as mean ± SEM.

### MSEW downregulates Gria2 mRNA and increases the Gria1/Gria2 ratio in the mPFC of male mice

We sought to identify the effect of MSEW on the gene expression of Gria1 and Gria2 and the possible relation between depression-like behaviour and the levels of these genes. For this purpose, we performed qPCR in the mPFC of males and females after the TST. For Gria1, we did not find significative differences (Figure 3A). Results for the Gria2 showed a significative main effect of *rearing* (F_1,36_=5.939, p<0.05) and the interaction *sex* × *rearing* (F_1,36_=10.577, p<0.01) (Figure 3A). The *rearing* effect showed that MSEW significantly decreased the transcription of Gria2 (p<0.05). The interaction *sex* × *rearing* showed a decreased Gria2 mRNA level in SN females in comparison with SN males (p<0.01). Moreover, the interaction also showed that maternally separated males had downregulation of Gria2 mRNA level in comparison with the SN males (p<0.001). For the Gria1/Gria2 ratio and the statistical analysis showed a significative difference for the interaction *sex* × *rearing* (F_1,36_=9.516, p<0.01) (Figure 3A). Bonferroni post-hoc test revealed a significant increased Gria1/Gria2 ratio (p<0.05) in MSEW males (p<0.01) compared to the SN males. Additionally, the analysis showed that MSEW females had a decreased Gria1/Gria2 ratio than the MSEW males (p<0.05).

**Figure 3.**
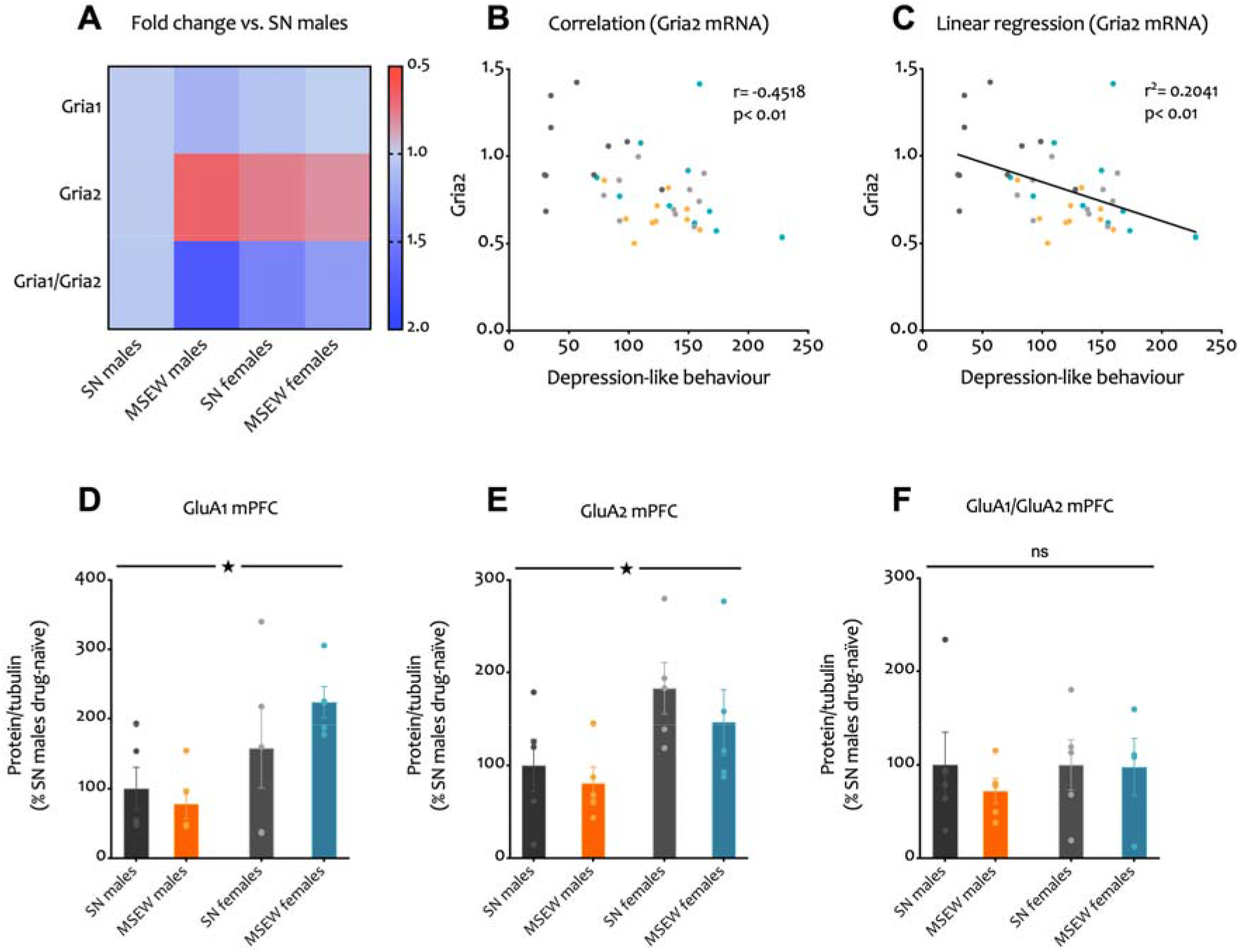
mRNA and protein levels of two AMPArs subunits in the mPFC of SN and MSEW drug-naïve mice. (A) Heatmap representing mRNA expression of Gria1, Gria2 and Gria1/Gria2 ratio in the mPFC of SN and MSEW mice relative to SN males as determined by qPCR. (B) Scatter blot illustrating the negative correlation between Gria2 and depression-like behaviour in mice. (C) Linear regression representing the relationship between depression-like behaviour and Gria2 expression in mice (Y= −0.0022X + 1.075). Mean fold change relative to SN males of (D) GluA1, (E) GluA2 and (F) GluA1/GluA2 protein levels in the mPFC of drug-naïve mice. *Sex* main effect of the ANOVA (⍰p<0.05). Data are expressed as mean ± SEM (n=5, run in duplicate or triplicate).

### Depressive-like behaviour correlates with Gria2 mRNA levels in the mPFC of mice

The relationship between change in Gria2 mRNA and the immobility time during the TST was determined using Pearson correlation coefficient. There was a significant correlation (r=−0.4518, p<0.01, n=40) between Gria2 and the depression-like behaviour among the subjects (Figure 3B). Linear regression suggests that Gria2 mRNA level could be predicted from the depression-like behaviour (F_1,38_=9.744, p<0.01) (y=−0.002244x+1.075) (Figure 3C).

### Increased GluA1 and GluA2 protein expression in female compared to male mice

Because of the differences observed in the Gria1 and Gria2 mRNA of mice, we decided to evaluate whether these changes were also manifested in the functional form of these genes. Hence, we detected the protein expression of GluA1 (Figure 3D), GluA2 (Figure 3E) and the GluA1/GluA2 ratio (Figure 3F), using western blotting in male and female mPFC of drug-naïve mice. The two-way ANOVA showed a main *sex* effect for GluA1 (F_1,16_=8.006, p<0.05) and GluA2 (F_1,16_=7.087, p<0.05). In both cases, the sex effect evidenced that females showed an increased protein expression compared to males. No changes were found for the GluA1/GluA2 ratio.

### Females show higher impulsive cocaine-seeking independently of the early-life stress

In order to determine if depressed animals exposed to MSEW showed different impulsivity for cocaine, we performed the cocaine SA procedure and calculated the average of the percentage of response efficiency along the acquisition phase (Figure 4A). The two-way ANOVA for the average of the response efficiency showed a main effect of *sex* (F_1,92_=8.016, p<0.01) and the interaction *sex* × *rearing* (F_1,92_=5.011, p<0.05). The *sex* effect indicated a lower response efficiency in female mice, meaning that females showed higher impulsivity to seek cocaine than males (p<0.01). The interaction *sex* × *rearing* revealed that maternally separated male mice had lower response efficiency than SN males (p<0.01), suggesting that MSEW increases impulsivity for cocaine-seeking but only in males. This interaction also evidenced that SN females showed higher cocaine-seeking impulsivity (decreased response efficiency) than SN males (p<0.01), behaving in the same way than the MSEW males and the MSEW females.

**Figure 4.**
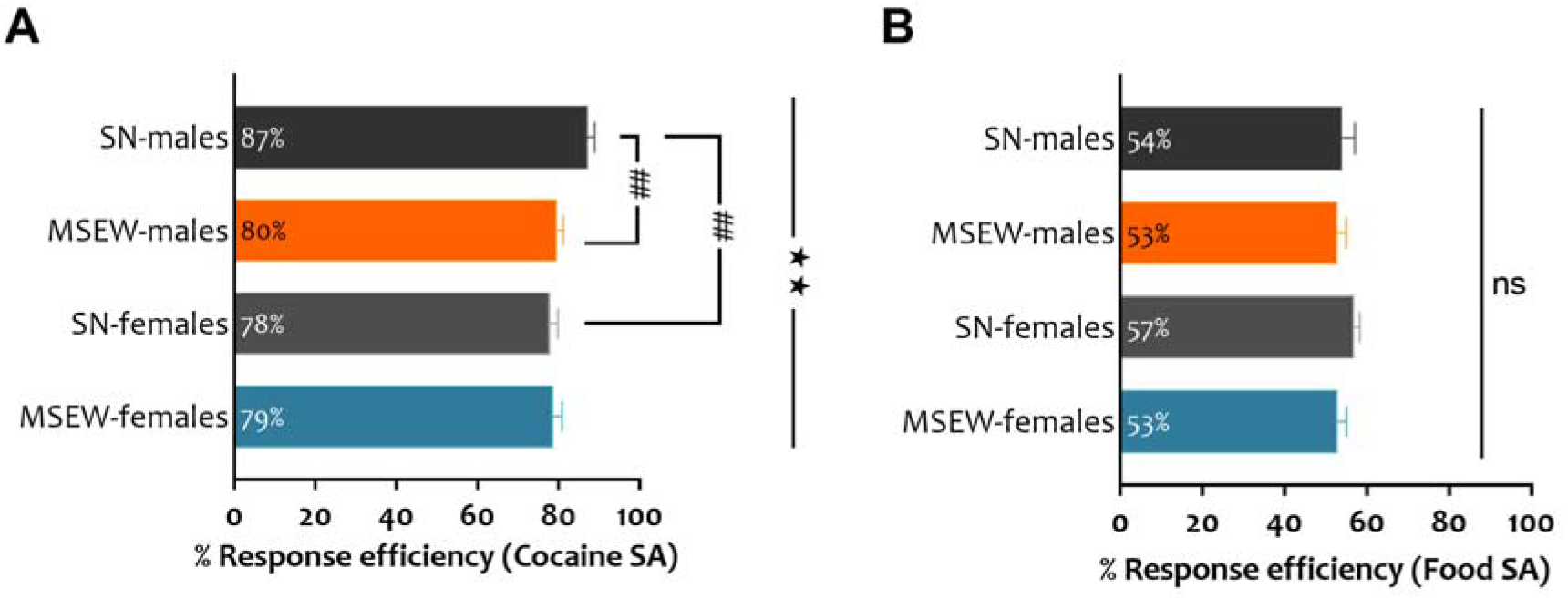
Percentage of response efficiency in the SA paradigm of SN and MSEW mice. Percentage of response efficiency as an indirect measure of impulsivity in the (A) cocaine SA or the (B) food SA procedure. *Sex* main effect of the ANOVA (⍰⍰p<0.01). Bonferroni post-hoc comparison for the interaction *sex* × *rearing* is indicated with the lines (##p<0.01). High response efficiency scores mean less impulsivity whereas low response efficiency signifies more impulsivity. Data are expressed as mean ± SEM.

### Cocaine but not natural reward SA is affected by early-life stress in male mice

To evaluate if the MSEW-induced increased impulsivity was specific for cocaine, we performed a food SA in another set of animals. Once again, we calculated the average of the percentage of response efficiency along the acquisition phase (Figure 4B). The two-way ANOVA did not show *sex* effect, *rearing* effect or the interaction between these two factors, suggesting that changes in impulsivity induced by MSEW affected cocaine-related impulsivity rather than responses related to natural reward.

### Chronic cocaine exposure does not modify the expression of GluA1, GluA2 or the GluA1/GluA2 ratio in the mPFC of mice

In order to evaluate changes in the expression of GluA1 (Figure 5A), GluA2 (Figure 5B) or the GluA1/GluA2 ratio (Figure 5C) due to cocaine exposure, we determined the levels of these proteins in the mPFC of mice that accomplish the cocaine SA. We also compared the values of these animals with the values of the drug-naïve mice. Three-way ANOVA revealed a *sex* effect for GluA1 (F_1,32_=31.094, p<0.001) and GluA2 (F_1,32_=6.905, p<0.05). In both cases, this main effect showed that females had higher protein expression than males. Moreover, statistical analysis showed a *sex* × *rearing* interaction for GluA1. The Bonferroni post-hoc test for the interaction specified that: SN females showed a higher GluA1 level than the SN males (p<0.05), MSEW females, higher than MSEW male mice (p<0.001) and also that MSEW females had increased protein expression than the SN females (p<0.05). For the GluA1/GluA2 ratio we did not find any significative differences.

**Figure 5.**
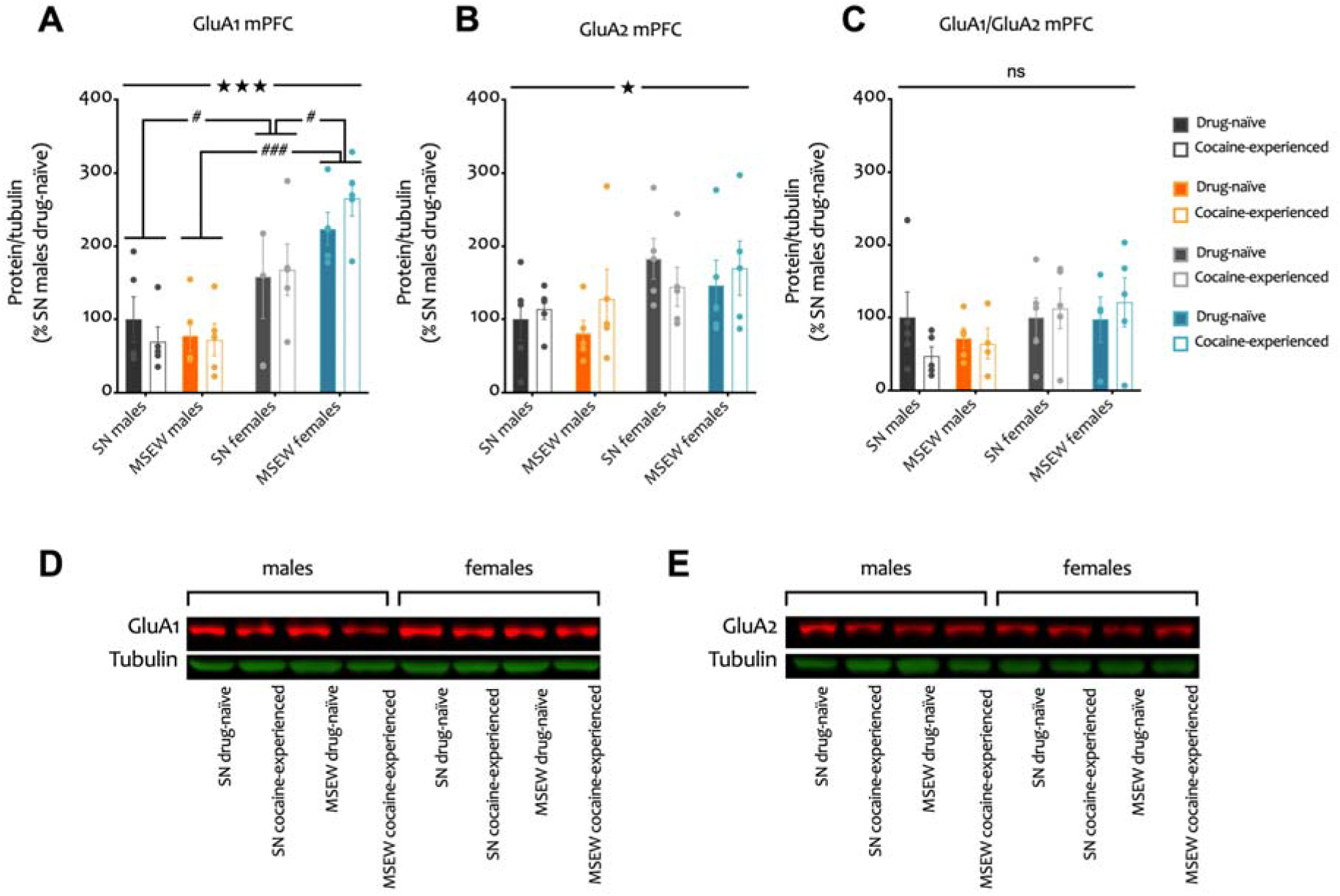
Fold change of GluA1, GluA2 and GluA1/GluA2 ratio in the mPFC of SN and MSEW mice. Mean fold change relative to SN males drug-naïve of (A) GluA1, (B) GluA2 and (C) GluA1/GluA2 levels in the mPFC. Representative western blot showing protein levels of (D) GluA1 and (E) GluA2 in the mPFC. The protein of interest in red and tubulin in green. *Sex* main effect of the ANOVA (⍰p<0.05, ⍰⍰⍰p<0.001). Bonferroni post-hoc comparison for the interaction *sex* × *rearing* (#p<0.05, ###p<0.001). Data are expressed as mean ± SEM (n=5, run in duplicate or triplicate).

## DISCUSSION

The present study shows that early-life stress induces a depressive-like behaviour in adult male mice whilst MSEW-exposed females seem to be resilient to this type of stress. Moreover, glutamatergic neurotransmission in the mPFC was differently affected due to rearing conditions and sex. Thus, Gria2 levels in the mPFC of MSEW male mice were different than in the SN males, showing that MSEW affected both sexes differently. However, in the case of GluA2, protein levels were not different between the SN males than to those exposed to early-life stress (MSEW). As measured in the cocaine SA, female mice showed increased impulsivity for cocaine-seeking independently of the early-life stress exposure. However, males were significantly affected by the MSEW, which increased their impulsivity for cocaine-seeking. Moreover, we observed that the MSEW-increased impulsivity in males was specific for cocaine because all the groups showed similar percentages of response efficiency in the food SA. Results from western blot in drug-naïve animals showed that female mice expressed higher levels of GluA1 and GluA2 in the mPFC, which could explain why they manifest higher impulsivity to cocaine intake and resilience to early-life stress. In sum, our results show a sex-dependent effect in the impulsivity to cocaine consumption and this effect seems to be cocaine-specific.

In previous work, we have also reported that only males were negatively affected by the MSEW in the acquisition of cocaine SA (Castro-Zavala *et al.* 2020a), showing increased excitability in the nucleus accumbens (increased GluA1/GluA2 ratio), while MSEW-exposed females did not express changes. In the present study, we do confirm that females show resilience to this type of stress.

The mPFC is a region that modulates cognitive and executive functions, including inhibitory control (Narayanan and Laubach 2017). Recent studies show that maternal separation in mice induces depression-like behaviours as well as a reduction of serotonin and dopamine levels in the frontal cortex (Récamier-Carballo *et al.* 2017). Additionally, animal models of depression suggest a reduced glutamate level in the PFC of depressed mice (Belin *et al.* 2008), as well as reduced glutamate and glutamate/glutamine levels in depressed rats (Li *et al.* 2008). Clinical evidence reported increased Gria2 mRNA levels in PFC in patients with major depression of both sexes, and no changes in Gria1 expression (Kleinman *et al.* 2015). Additionally, it was reported reduced glutamate/glutamine and GABA levels in the PFC of depressed patients (Hasler *et al.* 2007). These previous studies are in accordance with the fact that the glutamatergic system, especially in the frontal cortex, plays a key role in the modulation of the depressive phenotype.

Our present results showed a negative correlation between Gria2 in the mPFC and depression-like behaviour, meaning that a decreased Gria2 correlates with increased immobility time in the TST. Our biochemical analysis also showed a significative reduction in the Gria2 levels of mice exposed to MSEW of both sexes. Additionally, we observed a significant reduction in the Gria2 mRNA levels of MSEW-exposed male mice that evidences a significant mood alteration. However, we did not observe significant differences in GluA1 or GluA2 protein level in MSEW exposed mice. Our results are in accordance with Ganguly *et al.* (2019) who reported that males maternally separated showed decreased in mPFC-Gria2 expression compared to control males. Likewise, they observed that females in general, showed higher GluA2 protein level than male mice, similar to our findings.

In accordance with our results, a study using a model of depression in rats evidenced a dysregulation in glutamate neurotransmission but without changes in the levels of GluA1 or GluA2 in the mPFC of depressed rats (Treccani *et al.* 2016). Additionally, other authors reported no changes in Gria1 mRNA level in the mPFC of depressed mice (Belin *et al.* 2008) in agreement with our results.

Epidemiologic studies showed that childhood adversity increased the risk of depression by 28.4% but also to illicit drug use by 16.5% (Anda *et al.* 2006). Animal studies exploring how stressful situations influence addiction-like behaviours described that acute and chronic stress alters the phosphorylation of GluA2, affecting the function of the AMPA receptor but no the GluA1/GluA2 ratio in the nucleus accumbens and hippocampus (Ellis *et al.* 2017; Caudal *et al.* 2010; Caudal *et al.* 2016). This evidence could explain why we not able to find differences in the GluA1/GluA2 ratio. It is possible that MSEW-exposed male mice showed decreased-induced phosphorylation of GluA2, which did not modify the GluA2 levels, but altered indirectly the excitability of the AMPA receptor function, in accordance with our original hypothesis.

Caffino *et al.* (2015) report that a single cocaine exposure in adolescent rats can reduce the number of dendritic spines without any changes of GluA1 or GluA2 in the total mPFC homogenate. Other studies employing the cocaine SA paradigm reported that disruption of the GluA2 phosphorylation potentiated the acquisition of cocaine SA (Ellis *et al.* 2017). In the current work, the biochemical analysis yielded a *sex* × *rearing* interaction for the GluA1, but any alteration induced *per se* by the drug exposure (Figure 5A). However, these results indicate that MSEW females have increased GluA1 than SN females and MSEW males. The behavioural results for the cocaine SA evidenced that females, independently of the early-life stress exposure have increased impulsivity (less response efficiency).

Clinical studies evaluate the levels of glutamate in lumbar cerebrospinal fluid (CSF) in subjects diagnosed with personality disorder and healthy volunteers (Coccaro *et al.* 2013). They observed a direct correlation between CSF glutamate levels and impulsivity and/or aggression, confirming the key role-playing by glutamate in the regulation of impulsivity (Coccaro *et al.* 2013). Moreover, it was observed that cocaine-dependent patients showed a negative correlation between the activation of the frontoparietal network (network related with the inhibition control) and the dependence severity (Barrós-Loscertales *et al.* 2019). In the present study, we observed that female mice have higher GluA1 protein expression at basal level (Figure 3D), which could be interpreted as greater glutamatergic activity in the mPFC than males. This fact could explain why females showed higher impulsivity than males due to increased glutamatergic activity in this brain area that seems to be related to the impulsivity regulation. Our hypothesis is in line with the association between the positive correlation between impulsivity and the number of AMPArs observed in the post-mortem anterior cingulate cortex of alcoholics (Kärkkäinen *et al.* 2013).

Following our suggestion, it could be expected that MSEW males exposed to cocaine, show an increase in the levels of GluA1. However, as discussed above, there is a possibility that the differences in the response efficiency during the cocaine SA between the SN males and the MSEW male mice can be explained by changes in the phosphorylation of GluA2. In the case of the MSEW females, the increased basal level of GluA1 could be compensating the hyperphosphorylation of GluA2, avoiding further increases on impulsivity for cocaine-seeking due to MSEW. Our hypothesis is in accordance with previous studies showing that methylphenidate enhances the response inhibition in rats because of the increased expression of AMPArs in the PFC (Zhang *et al.* 2017); that increased AMPArs contributes to modulate the activity of the NMDA receptor in order to learn to inhibit a response (Hayton *et al.* 2010).

Our results are also in line with recent findings in which mutant mice lacking GRIP1 in the mPFC, a scaffolding protein that stabilizes GluA2 at the surface, showed increased GluA2-containing AMPArs in the cell membrane, and increased cocaine intake during the SA paradigm (Wickens *et al.* 2019). Moreover, they observed that these effects were cocaine specific and GRIP1 does not influence natural reward-seeking, as we observed in the current work.

Taken together, our results propose that female mice exhibit increased basal mPFC glutamatergic function, which potentiates impulsivity to cocaine consumption during the SA. Additionally, MSEW alters the functionality of the glutamatergic circuits in male mice, increasing the glutamatergic excitability (GluA1/GluA2) in an indirect way. We also propose that MSEW-exposed males showed higher GluA2-containing AMPArs due to the altered GluA2, which avoid the internalization of this subunit. Additionally, we suggest that early-life stress affects several molecular mechanisms avoiding the ability to stabilize GluA2 in the synaptic surface. Our study evidences the underlying molecular mechanisms of the glutamatergic system regulating impulse control disorders, like cocaine use disorder and their relationship with the depression phenotype.

In addition, the present study provides novel evidence regarding the molecular mechanisms altered in the glutamatergic system in the prefrontal cortex of mice exposed to early-life stress during the first postnatal days. These alterations could underlie the higher cocaine motivation due to modifications in the inhibition control.

## Acknowledgements

This study was supported by the Ministerio de Economia y Competitividad (grant number SAF2016-75966-R-FEDER), Ministerio de Sanidad (Retic-ISCIII, RD16/017/010 and Plan Nacional sobre Drogas 2018/007) to O.V., and the Consejo Nacional de Ciencia y Tecnología in Mexico (CONACYT) (CB-2015-255317 to A.C., 582196 for L.M.-M. and 276577 for A.C-Z). The authors thank Inés Gallego-Landin for her English proofreading and editing of the manuscript.

## Author contributions

A.C-Z. and O.V. were responsible for the study concept and design. A.C-Z., A.M-S. and L.M-M. carried out the experimental studies. A.C-Z and O.V. drafted the manuscript and A.C-Z, O.V and A.C.-M. participated in the interpretation of findings. All authors critically reviewed the content and approved the final version for publication.

## ABBREVIATIONS

AMPArs: α-amino-3-hydroxy-5-methyl-4-isoxazolepropionic acid receptors
CSF: Lumbar cerebrospinal fluid
GABA: Gamma-aminobutyric acid
GluA1: AMPA receptor subunit 1
GluA2: AMPA receptor subunit 2
Gria1: Glutamate Ionotropic Receptor AMPA Type Subunit 1 gen
Gria2: Glutamate Ionotropic Receptor AMPA Type Subunit 2 gen
MDD: Major depression disorder
NAc: Nucleus accumbens
mPFC: medial prefrontal cortex
MSEW: Maternal separation with early weaning
PD: Postnatal day
PFC: Prefrontal cortex
SA: Self-administration
SN: Standard nest
SUD: Substance use disorder
TST: Tail suspension test
VTA: Ventral tegmental area

